# Horizontal transfer of a pathway for coumarate catabolism unexpectedly inhibits purine nucleotide biosynthesis

**DOI:** 10.1101/315036

**Authors:** Dan M. Close, Sarah J. Cooper, Xingyou Wang, Richard J. Giannone, Nancy Engle, Timothy J. Tschaplinski, Lizbeth Hedstrom, Jerry M. Parks, Joshua K. Michener

## Abstract

Metabolic pathways are frequently transferred between bacterial strains in the environment through horizontal gene transfer (HGT), yet laboratory engineering to introduce new metabolic pathways often fails. Successful use of a pathway requires co-evolution of both pathway and host, and these interactions may be disrupted upon transfer to a new host. Here we show that two different pathways for catabolism of coumarate failed to function when initially transferred into *Escherichia coli*. Using laboratory evolution, we elucidated the factors limiting activity of the newly-acquired pathways and the modifications required to overcome these limitations. Both pathways required mutations to the host to enable effective growth with coumarate, but the necessary mutations differed depending on the chemistry and intermediates of the pathways. In one case, an intermediate inhibited purine nucleotide biosynthesis, and this inhibition was relieved by single amino acid mutations to IMP dehydrogenase. A strain that natively contains this coumarate catabolism pathway, *Acinetobacter baumannii*, is already resistant to inhibition by the relevant intermediate, suggesting that natural pathway transfers have faced and overcome similar challenges. These discoveries will aid in our understanding of HGT and ability to predictably engineer metabolism.

This manuscript has been authored by UT-Battelle, LLC under Contract No. DE-AC05-00OR22725 with the U.S. Department of Energy. The United States Government retains and the publisher, by accepting the article for publication, acknowledges that the United States Government retains a non-exclusive, paid-up, irrevocable, world-wide license to publish or reproduce the published form of this manuscript, or allow others to do so, for United States Government purposes. The Department of Energy will provide public access to these results of federally sponsored research in accordance with the DOE Public Access Plan (http://energy.gov/downloads/doe-public-access-plan).

## Introduction

Microbes have the ability to use a wide variety of compounds as carbon and energy sources. Expanding the breadth of compounds that a strain can catabolize can be highly beneficial, both to access to new environmental niches and for engineered microbes that can use new feedstocks. Correspondingly, the catabolic pathways responsible for these abilities are frequently transferred between strains, either in nature through horizontal gene transfer (HGT) or in the laboratory through metabolic engineering (Nielsen and Keasling, 2016; Pál et al., 2005). However, depending on the precise chemistry involved, new pathways often fail to function effectively in their new host (Porse et al., 2018). In these cases, productive use of a new pathway may require post-transfer refinement to optimize expression and minimize deleterious interactions (Clark et al., 2015; Michener et al., 2014). The pathway activity immediately following transfer may be very different from the potential activity after optimization, complicating predictions about engineering or HGT.

Pathway selection and optimization is particularly important for the conversion of lignin-derived aromatic compounds, such as the phenylpropanoid coumarate. Use of lignocellulosic biomass as a feedstock for biofuel production yields a substantial lignin byproduct stream, which can be thermochemically depolymerized to yield complex mixtures containing multiple phenylpropanoid derivatives (Rodriguez et al., 2017). Many of these phenylpropanoids also occur naturally during lignocellulose decay and, as a result, microbes have evolved the ability to consume them as sources of carbon and energy (Bomble et al., 2017; Bugg et al., 2011). The efficient biological conversion of lignin-derived aromatic compounds into fuels and chemicals will be a key factor in making biofuel production cost-effective (Linger et al., 2014; Ragauskas et al., 2014). Successful use of the diverse mixtures of compounds produced from lignin depolymerization will require the ability to tailor a metabolic network to the particular substrate mixture. Facile assembly of a metabolic network from individual pathways will, in turn, require the elimination of any inhibitory interactions between pathways and the host.

We have explored these issues using pathways for catabolism of a natural phenylpropanoid, coumarate, transferred into the non-native host *E. coli*. There are two known oxidative routes for phenylpropanoid catabolism, differing in their specific reaction chemistry and resulting intermediates (Figure 1). These pathways are exemplified by the *hca* pathway from *Acinetobacter* sp. ADP1 (Parke and Ornston, 2003) and the *cou* pathway from *Rhodococcus jostii* (Otani et al., 2014). Both pathways begin by deacetylating the phenylpropanoid substrate. The *hca* pathway then uses a retro-aldol reaction to produce an intermediate benzaldehyde derivative, while the *cou* pathway uses a hydrolytic retro-Claisen reaction to directly produce the benzoate derivative.

**Figure 1:**
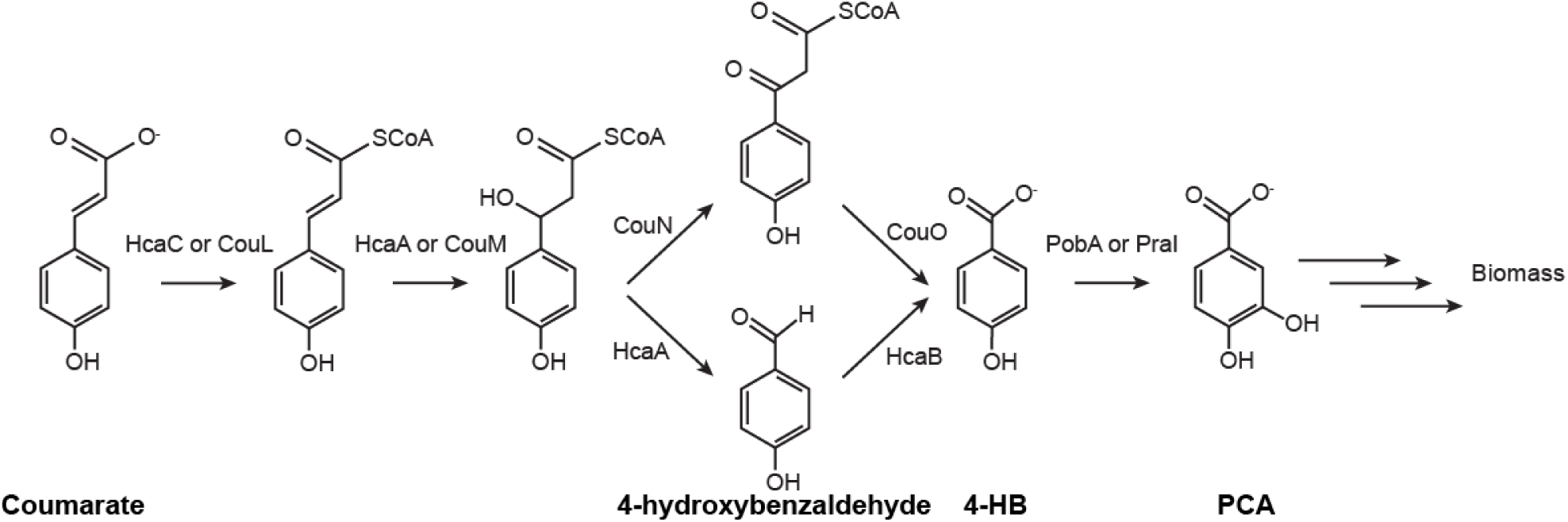
Two routes, exemplified by the *hcaABC* pathway from *Acinetobacter* sp. ADP1 and the *couLMNO* pathway from *R. jostii*, deacetylate the phenylpropanoid coumarate to 4-hydroxybenzoate. For simplicity, cofactors and the resulting acetyl-CoA are not shown.

However, biochemical pathways do not function in isolation, but instead are embedded in a complex network of metabolic and regulatory interactions. Transfer into a new host will disrupt these interactions, potentially interfering with either activity of the heterologous pathway or the host’s native processes. Since these two phenylpropanoid pathways use different biochemistry and intermediates, their interactions with the host may also differ substantially (Kim and Copley, 2012). Identifying the likeliest pairing of host and pathway, either for engineering or HGT, will depend on understanding the specific challenges imposed by each potential pathway and the mechanisms to overcome these challenges available to the host.

In this work, we used a combination of engineering and evolution to construct and optimize two pathways for phenylpropanoid catabolism in the common host, *E. coli*. We show that, after optimization, both the *hca* and *cou* pathways are capable of supporting growth with coumarate. However, due to differences in pathway biochemistry, the mutations required for efficient growth differ between the two pathways. The *hca* pathway produces a unique metabolite that inhibits a key enzyme in nucleotide biosynthesis, and mutations to the host are necessary to alleviate this inhibition and allow growth. Beneficial mutations to the *cou* pathway occurred through different mechanisms, including mutations to an enzyme involved in cofactor salvage. Understanding these types of interactions between an engineered metabolic pathway and its heterologous host is key to building flexible metabolic networks that can easily be tailored to specific feedstocks.

## Results

### Combining engineering and evolution enabled coumarate catabolism

We designed and synthesized two pathways for phenylpropanoid import and degradation, each of which converts coumarate into 4-hydroxybenzoate (4-HB) (Figure 1 and Figure S1). Each pathway was introduced into *E. coli* strains, JME38 and JME50, that had previously been engineered to grow with 4-HB (Standaert et al., 2018). None of the engineered strains acquired the immediate ability to grow with coumarate as the sole source of carbon and energy (Figure S2).

To understand the factors preventing pathway function, we used experimental Three replicate cultures of each engineered strain were propagated in minimal medium containing 1 g/L coumarate. After 300 generations, individual mutants were isolated from each population and characterized for growth with protocatechuate (PCA), 4-HB, coumarate, and caffeate. Representative isolates were chosen for each replicate population for further characterization. All isolates could grow with PCA and coumarate, though growth with caffeate and 4-HB varied between replicates (Figure S2).

### Genome resequencing and reconstruction identified causal mutations

The genomes of the selected isolates were resequenced to identify new mutations (Table S1). Several of the mutations have previously been described for their effects on catabolism of 4-HB, such as silent mutations to the gene encoding the 4-hydroxybenzoate monooxygenase *pobA* (Standaert et al., 2018). Among the strains with the *hca* pathway, five of the six isolates had additional mutations to the native gene *guaB*, encoding inosine monophosphate (IMP) dehydrogenase (IMPDH), and to the intergenic region between *hcaB* and *hcaC* in the engineered pathway. The exception was JME96, which had a mutation to *rpoS* that is expected to be highly pleiotropic (Saxer et al., 2014).

In the strains with the *cou* pathway, the acquired mutations were less consistent across replicates, with several mutations to genes that are expected to be pleiotropic. However, parallel mutations were observed in JME106 and JME109, with mutations to both *couL* and *nadR*. The mutations to *couL*, which encodes the CoA ligase, were coding mutations, L192R and S134Y. One of the mutations to *nadR* led to a frameshift that precisely removed the C-terminal ribosylnicotinamide kinase (RNK) domain and the second *nadR* mutation also occurred in the RNK domain (Kurnasov et al., 2002).

To test the causality of the identified mutations, we reconstructed representative mutations in the engineered parental strains. Two mutations, to *pobA* and *hcaABCK,* were necessary for growth with coumarate in JME64, while a third mutation to *guaB* significantly increased growth (Figure 2A). Similarly, mutations to *pobA*, *couLHTMNO*, and *nadR* were all required for growth with coumarate using the *cou* pathway in JME65 (Figure 2B).

**Figure 2:**
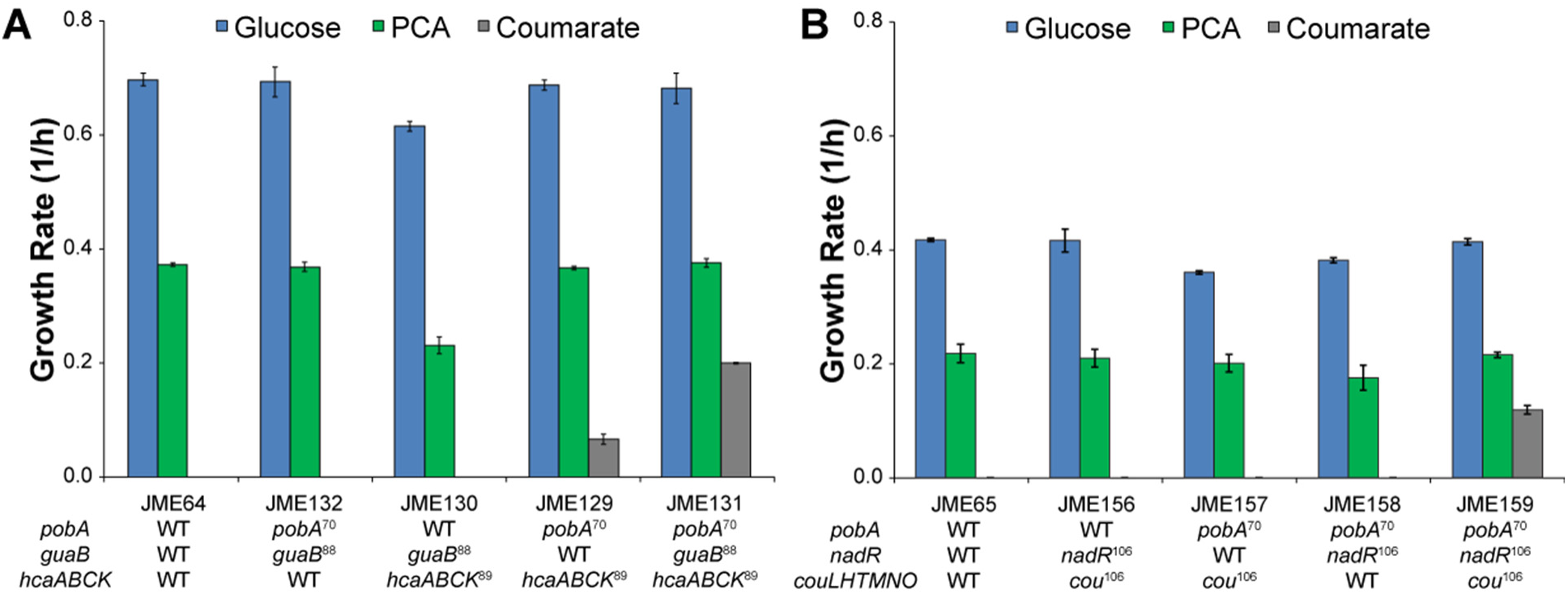
Reconstruction identifies causal mutations. (A) For the *hca* pathway, a triple mutant, containing mutations to *pobA*, *guaB*, and *hcaABCK*, and the three double mutants were grown in minimal medium with 1 g/L of the indicated substrate as the sole source of carbon and energy. (B) As in A, except using the *cou* pathway and mutations to *pobA*, *nadR*, and *couHLTMNO*. Error bars show one standard deviation, calculated from three biological replicates. For complex mutations, allele superscripts indicate the evolved strain from which that allele was taken.

The *pobA* mutation has previously been shown to increase expression of PobA by destabilizing secondary structures in the mRNA (Standaert et al., 2018). To understand the effect of the intergenic mutations upstream of *hcaC*, we measured protein expression levels in the engineered strains. As expected, the mutation to *pobA* increased expression of PobA by approximately 9-fold, while the *hcaC* mutation increased expression of both HcaB and HcaC by roughly 2-fold (Figure S3A). The intergenic mutation before *hcaC* is predicted to increase the translation rates by approximately 10-fold (Espah Borujeni et al., 2014).

Parallelism of mutations within replicates of a pathway, but divergence between pathways, strongly suggests that the mutations are specific to a particular pathway. To test this hypothesis, we replaced the *hca* pathway in JME131 with either the wild-type or evolved *cou* pathways. Neither strain was able to growth with coumarate as the sole source of carbon and energy.

### Inhibitory cross-talk between engineered and native pathways

A mutation to *guaB* was necessary for growth with coumarate using the *hca* pathway. IMPDH, encoded by *guaB*, converts inosine monophosphate (IMP) to xanthosine monophosphate (XMP) during purine nucleotide biosynthesis (Hedstrom, 2009). Five independent mutations to IMPDH were identified: A48V, D243G, G330D, L364Q, and P482L. We generated a model of the *E. coli* IMPDH, using multiple IMPDH crystal structures as templates and mapped the positions of mutations onto the model (Figure 3A). The mutations are scattered around the structure, with no evident common mechanism to affect the active site (Figure 3B).

**Figure 3:**
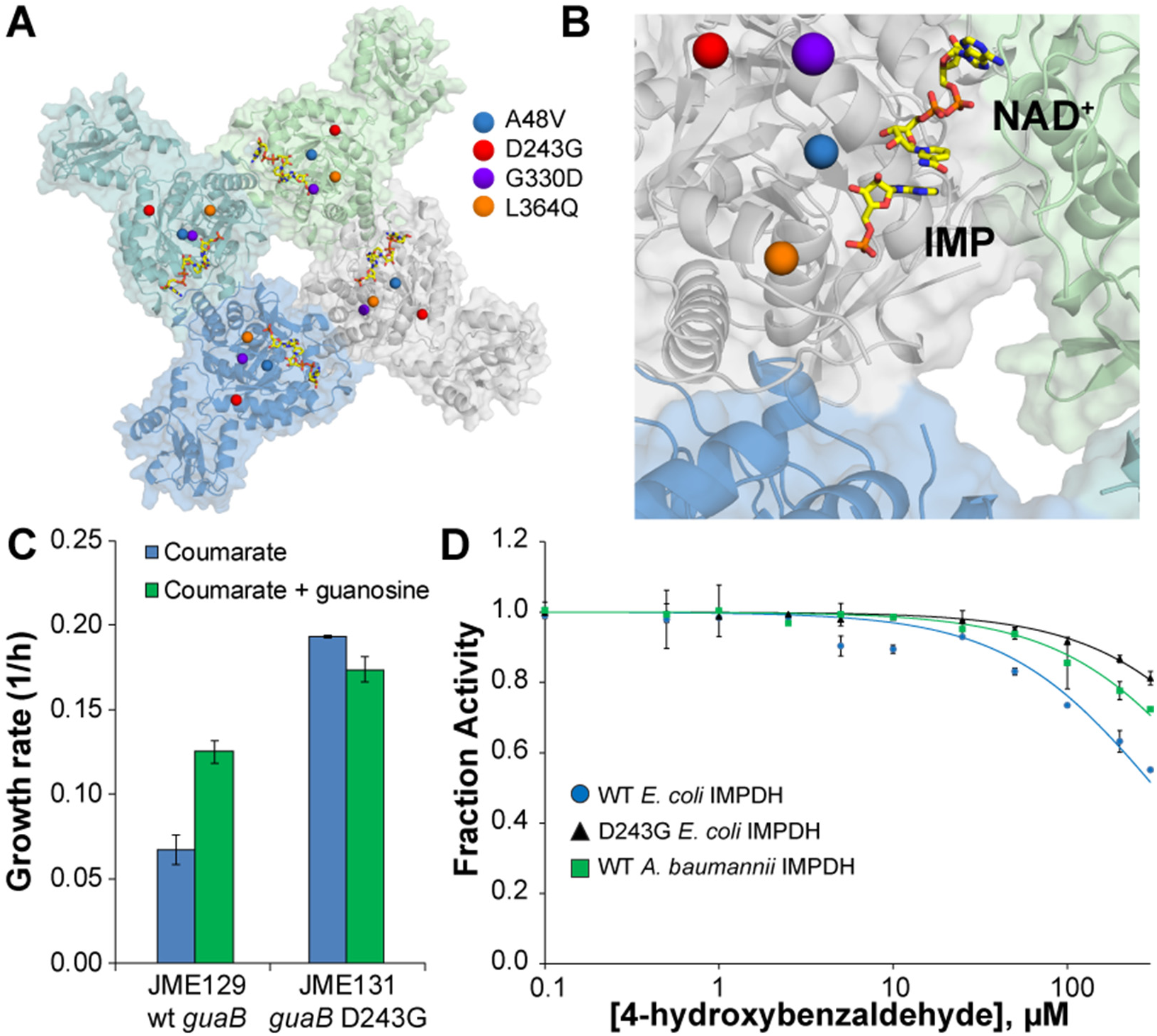
Accumulation of 4-hydroxybenzaldehyde inhibits IMPDH. (A) Positions of mutated residues in a model of the *E. coli* IMPDH. P482 was not included in the model and is not shown. Each subunit of the IMPDH tetramer is shown in a different color. (B) A detailed view of the active side of the model. IMP and NAD^+^ are shown docked in the active site. Mutated residues are as in A. (C) Strains with wildtype and mutant versions of IMPDH were grown in medium containing 1 g/L coumarate with and without the addition of 5 mg/L guanosine. No growth was seen with guanosine alone. Error bars show the standard deviation, calculated from three biological replicates. (D) Enzyme variants were purified and assayed *in vitro* for inhibition by 4-hydroxybenzaldehyde. Curves show a model fit, using the calculated inhibition constants. Error bars show the standard deviation, calculated from three biological replicates.

To understand the consequences of these mutations, we measured metabolite levels in the parent and engineered strains during growth with coumarate. Compared to the D243G *guaB* mutant, the strain with wild-type *guaB* showed higher levels of AMP (Figure S4). Concentrations of GMP and IMP were below the limit of detection of our assay. These results suggested that growth with coumarate perturbed the purine nucleotide pools.

We hypothesized that growth with coumarate led to inhibition of IMPDH and depletion of guanine nucleotides, and that this inhibition was relieved in the *guaB* mutants. To determine whether guanine nucleotide depletion inhibited growth with coumarate, we supplemented the growth medium with guanosine. Addition of guanosine increased growth with coumarate in a strain with the wildtype IMPDH, but not the mutant (Figure 3C).

Mutations to IMPDH improved growth with the *hca* pathway but not with the *cou* pathway. The *hca* pathway produces an intermediate, 4-hydroxybenzaldehyde, that is not present in the *cou* pathway (Figure 1). To test whether this intermediate was responsible for the inhibition of IMPDH, we purified WT and mutant IMPDH and measured inhibition *in vitro* with 4-hydroxybenzaldehyde. This compound is a weak inhibitor of WT *Ec*IMPDH, with a *K*_i,app_ of 320 ± 20 μM (Figure 3C). Introduction of the D243G mutation had little effect on catalytic activity (Table S5) but increased the *K*_i,app_ to 1250 ± 50 μM, indicating a substantial reduction of inhibition in the mutant. The *hca* pathway that we used came from *Acinetobacter* sp. ADP1, and we hypothesized that the native IMPDH of this strain would have faced similar selective pressures to minimize inhibition by 4-hydroxybenzaldehyde. As a surrogate, we tested the IMPDH of *A. baumanii*, which contains a homologous *hca* pathway. As predicted, the *A. baumannii* IMPDH has a *K*_i,app_ of 720 ± 30 μM. Further kinetic characterization of these IMPDH homologs is summarized in Table S5.

In the IMPDH model, the carboxylate side chain of D243 forms hydrogen bonds with the backbone of V220 and the side chain of K87, which would be disrupted in the D243G mutant (Figure S6). K87 is located at the C-terminal end of a long α helix, and V220 is at the beginning of a β strand. It is not obvious how these local changes might be propagated to the active site to relieve inhibition without affecting catalysis.

To gain insight into possible mechanisms of inhibition by 4-hydroxybenzaldehyde, we computationally docked 4-hydroxybenzaldehyde to the wild-type IMPDH model in the apoenzyme, IMP-bound, and IMP/NAD^+^-bound states (Figure 4). As a test of our modeling and docking approach, we first redocked IMP and NAD^+^ into their respective binding sites in the apoenzyme and compared the resulting models with relevant template structures containing these molecules. Whereas the top docked pose of NAD^+^ deviates slightly (1.1 Å RMSD) from its position in the crystal structure, IMP is essentially superimposable (0.3 Å RMSD) with the corresponding crystallographic coordinates of XMP (PDB entry 4X3Z). Thus, we deemed our approach sufficiently accurate to dock 4-hydroxybenzaldehyde to each of the three models. In both the apoenzyme and IMP-bound models, the majority of the top poses of 4-hydroxybenzaldehyde occupy the NAD^+^ binding site (Table S7). However, the structural changes that we predict would occur in the mutant enzymes do not significantly change these binding interactions.

**Figure 4.**
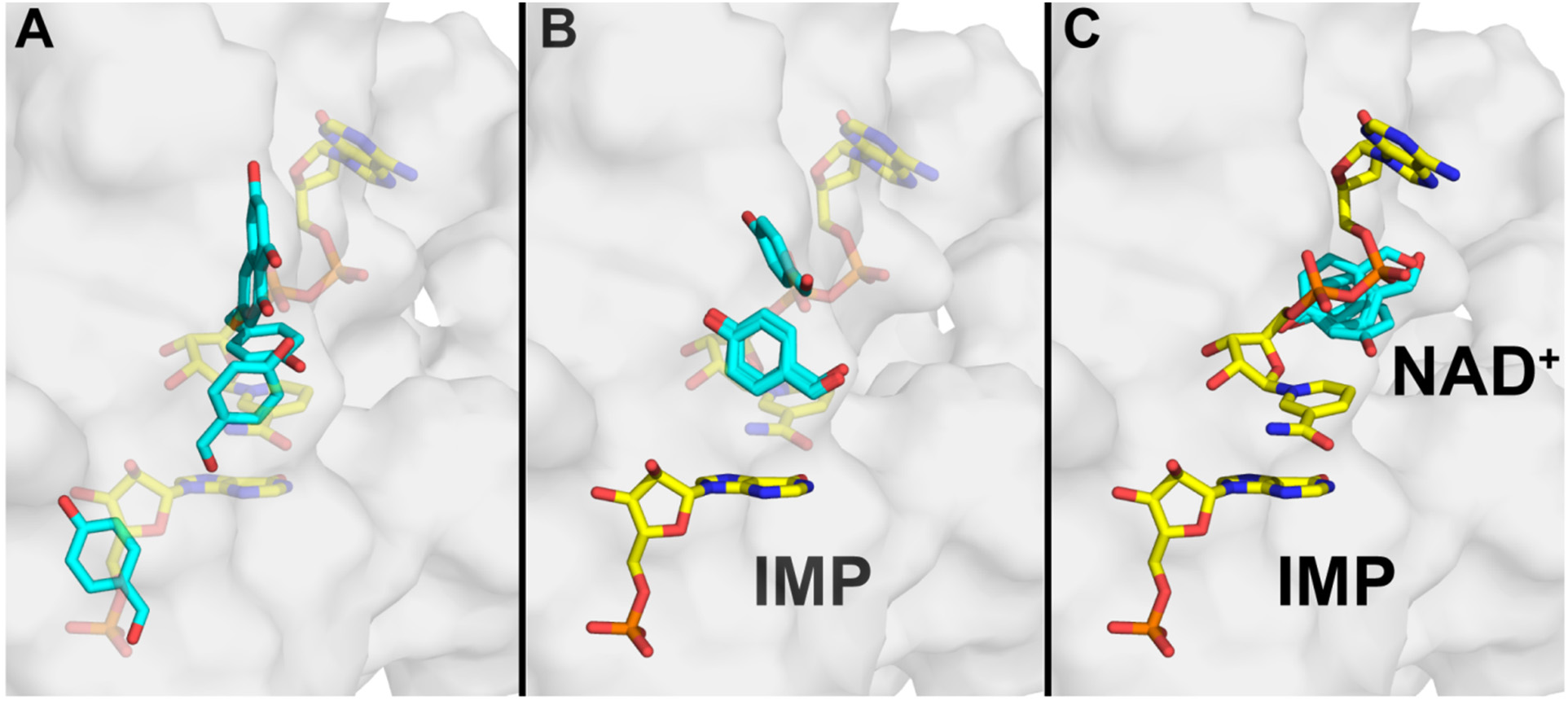
Predicted docking poses for 4-hydroxybenzaldehyde to IMPDH in various enzyme states. (A) Apoenzyme, (B) IMP-bound, and (C) IMP/NAD^+^-bound IMPDH. The top five docking poses are shown in each case. 4-hydroxybenzaldehyde carbons are shown in cyan. All other carbons are shown in yellow. Molecules in transparent representation are shown for reference, but were not included in the docking. All hydrogens are omitted for clarity.

## Discussion

The *hca* pathway shows clear signs of HGT, with highly homologous pathways present in various beta- and gamma-proteobacteria. In this work, we have recapitulated the process of HGT, and demonstrated the necessity for host adaptations to accommodate the pathway in both *E. coli* and *A. baumannii*. Further HGT of this pathway would require either a host with an IMPDH homolog that is resistant to inhibition by 4-hydroxybenzaldehyde, or post-transfer selection for mutations that relieve inhibition. Understanding these types of limitations on HGT, and the mechanisms by which organisms evolve to avoid them, will aid in our ability to predict and manipulate horizontal gene transfer (Clark et al., 2015; Michener et al., 2014).

In combination, our results suggest that introduction of the *hca* pathway did not allow growth with coumarate because accumulation of 4-hydroxybenzaldehyde inhibited the native *E. coli* IMPDH. This inhibitory cross-talk results in nucleotide starvation and impairs growth and phenylpropanoid catabolism. Mutations to IMPDH prevent inhibition by 4-hydroxybenzaldehyde and allow growth with coumarate. There is no *a priori* reason to expect that a pathway for degradation of an aromatic compound would interact with a native pathway for nucleotide biosynthesis. Phenolic amides such as feruloyl amide have been shown to inhibit a different step in nucleotide biosynthesis (Pisithkul et al., 2015), but neither the substrate nor products of coumarate degradation are toxic at the relevant concentrations (Clarkson et al., 2017; Standaert et al., 2018). These types of inhibitory cross-talk are likely to be common with introduced metabolic pathways, though they are rarely identified and alleviated (Kim and Copley, 2012; Kizer et al., 2008; Michener et al., 2012). As we have shown, relatively subtle changes in pathway structure, such as the differences between the *hca* and *cou* pathways, can dramatically change the interactions between a pathway and its host.

Across the replicate populations, many mutations were highly pleiotropic, including large insertions and deletions as well as mutations to core transcriptional machinery such as *rho* and *rpoB*. Duplications frequently spanned the insertion sites for engineered operons, suggesting that expression of the heterologous genes was limiting. By comparing across replicates, however, we were able to identify a set of point mutations that allowed growth with coumarate as the sole source of carbon and energy. However, a reconstructed strain containing these mutations does not grow as quickly with coumarate as the evolved isolates, suggesting that some of the remaining mutations provided additional fitness benefits (Figure 2 and Figure S2).

The phenylpropanoid CoA ligases, *couL* and *hcaC*, both required mutations for full heterologous activity. The mutations to *hcaC* increased expression, presumably by modulating translation, while the mutations to *couL* decreased expression (Figure S3B). Without direct measurements, we cannot say whether the mutations to CouL also affected the specific activity of the enzyme, or whether the observed change in expression is the sole explanation for the observed benefit. Similarly, it is unclear what role the *nadR* truncation plays in coumarate degradation using the *cou* pathway. The C-terminal ribosylnicotinamide kinase domain is involved in a minor pathway for NAD^+^ salvage. This domain may be promiscuously phosphorylating an intermediate in the *cou* pathway.

Inhibition of microbial growth by aldehydes is commonly observed, though the mechanisms of toxicity can rarely be traced to a specific interaction (Clarkson et al., 2014; Mills et al., 2009; Yi et al., 2015). Mutations that increase tolerance generally do so either by increasing export of the toxic compound or by performing redox chemistry to remove the aldehyde functionality (Mukhopadhyay, 2015). In this work, we have shown an example of aldehyde toxicity that acts through a single protein and can be relieved by point mutations to that protein. Other examples of nonspecific toxicity may prove to be similarly specific when characterized fully.

We have described the use of experimental evolution to identify and alleviate deleterious interactions between engineered metabolic pathways for coumarate catabolism and native pathways for nucleotide biosynthesis and cofactor salvage. Many engineered pathways place a substantial burden on the production host, but understanding and accommodating these interactions remains challenging. Evolution can simplify this optimization process by directly selecting for mutations that eliminate the inhibition. As we did with *guaB*, researchers can then work backwards from the evolutionary solutions to understand the factors that were initially limiting productivity and the biochemical solutions to overcome those problems. By solving more problems of this sort, we will develop design rules for future forward engineering of decoupled metabolic pathways and better predictions of the likelihood of pathway transfer by HGT.

## Materials and Methods

### Strains and chemicals

Unless otherwise noted, all chemicals were purchased from Sigma-Aldrich (St. Louis, MO) or Fisher Scientific (Fairlawn, NJ) and were molecular grade. All oligonucleotides were ordered from IDT (Coralville, IA). *E. coli* strains were routinely cultivated at 37 °C in LB containing the necessary antibiotics (50 mg/L kanamycin or 50 mg/L spectinomycin). Growth assays with aromatic substrates were performed in M9 salts medium containing 300 mg/L thiamine and 1 mM isopropyl β-D-1-thiogalactopyranoside (IPTG). PCA and 4-HB were dissolved in water at 5 g/L, filter sterilized, and added at a final concentration of 1 g/L. Coumarate and caffeate were dissolved in DMSO at 100 g/L and added at a final concentration of 1 g/L. The addition of 1% DMSO did not affect growth. The pH of the substrates was not controlled, as PCA oxidation occurred more rapidly at neutral pH.

### Plasmid construction

Plasmids pJM219 and pJM223, containing the *cou* and *hca* expression constructs, were synthesized by the Joint Genome Institute. As described previously, the pathway design used synthetic promoters, terminators, and custom ribosome binding sites (Chen et al., 2013; Espah Borujeni et al., 2014; Kosuri et al., 2013; Salis et al., 2009). Plasmids expressing sgRNA for chromosomal modifications were constructed as described previously, using an inverse PCR to linearize the expression vector followed by assembly with synthesized oligonucleotides (Clarkson et al., 2017). Plasmid pJM303, expressing the D243G mutant of the *E. coli* IMPDH, was constructed by amplifying the mutant *guaB* allele from JME89 and cloning it into pMCSG7 under the control of a T7 promoter.

### Strain construction

Genome modifications were performed as described previously, using the lambda-red recombineering system in combination with Cas9-mediated selection (Clarkson et al., 2017; Jiang et al., 2015). Integration cassettes were amplified from synthesized plasmids or the chromosomal DNA of mutant strains, as needed.

### Experimental evolution

Parental strains were streaked to single colonies. Three colonies from each strain were grown to saturation in LB + 1 mM IPTG, then diluted 128-fold into M9 + 1 mM IPTG + 1 g/L coumarate + 50 mg/L PCA and grown at 37 °C. When the cultures reached saturation, typically after two days during the initial stages, they were diluted 128-fold into fresh medium. As the growth became more robust, the PCA concentration was decreased. Cultures derived from JME65 and JME67 required the addition of PCA for 150 generations, and in some cases took two days to reach saturation even after 300 generations. One replicate culture, JME65-C, became contaminated with a different coumarate-degrading strain. This contamination was not discovered until resequencing, and consequently the culture was not restarted.

After 300 generations, the evolved cultures were streaked to single colonies. Six isolates from each replicate culture were tested for growth with PCA, 4-HB, coumarate, and caffeate. Representative isolates were selected and validated, followed by genome resequencing.

### Genome resequencing

Genomic DNA was isolated using a Blood and Tissue kit (Qiagen, Valencia, CA), according to the manufacturer’s directions. The DNA was then sequenced by the Joint Genome Institute on a MiSeq (Illumina, San Diego, CA) to approximately 75x coverage.

### Growth rate measurements

Growth rates were measured as described previously (Clarkson et al., 2017). Briefly, cultures were grown overnight to saturation in M9 + 1 mM IPTG + 2 g/L glucose. They were then diluted 100-fold into fresh M9 + IPTG containing the appropriate carbon source and grown as triplicate 100 μL cultures in a Bioscreen C plate reader (Oy Growth Curves Ab Ltd, Helsinki, Finland). Growth rates were calculated using CurveFitter software based on readings of optical density at 600 nm (Delaney et al., 2013).

### Proteomic measurements

Engineered *E. coli* strains were grown to saturation in 5 mL cultures of M9 + 2 g/L glucose + 1 mM IPTG. They were then diluted 100-fold into triplicate 5 mL of the same medium and grown to mid-log phase. The cells were separated by centrifugation, washed twice with water, and frozen in LN_2_ for later analysis.

Processing for LC-MS/MS analysis was performed as previously described (Clarkson et al., 2017). Briefly, crude protein lysates were obtained by bead beating cells in sodium deoxycholate (SDC) lysis buffer (4% SDC, 100 mM ammonium bicarbonate, pH 8.0). Cleared protein lysates were then adjusted to 10 mM dithiothreitol and incubated at 95 °C for 10 min to denature and reduce proteins. Cysteines were alkylated/blocked with 30 mM iodoacetamide and 250 μg transferred to a 10-kDa MWCO spin filter (Vivaspin 500, Sartorius) for *in situ* clean-up and digestion with sequencing-grade trypsin (G-Biosciences). The tryptic peptide solution was then spin-filtered through the MWCO membrane, adjusted to 1% formic acid to precipitate residual SDC, and SDC precipitate removed from the peptide solution with water-saturated ethyl acetate extraction. Peptide samples were then concentrated via SpeedVac (Thermo Fisher) and quantified by BCA assay (Pierce) prior to LC-MS/MS analysis.

Peptide samples were analyzed by automated 2D LC-MS/MS analysis using a Vanquish UHPLC plumbed directly in-line with a Q Exactive Plus mass spectrometer (Thermo Scientific) outfitted with a triphasic MudPIT back column (RP-SCX-RP) coupled to an in-house pulled nanospray emitter packed with 30 cm of 5 μm Kinetex C18 RP resin (Phenomenex). For each sample, 5 μg of peptides were loaded, desalted, separated and analyzed across two successive salt cuts of ammonium acetate (50 mM and 500 mM), each followed by 105 min organic gradient, as previously detailed (Clarkson et al., 2017). Eluting peptides were measured and sequenced by data-dependent acquisition on the Q Exactive MS.

MS/MS spectra were searched against the *E. coli* K-12 proteome concatenated with exogenous Pca, Hca, and Cou pathway proteins, common protein contaminants, and decoy sequences using MyriMatch v.2.2 (Tabb et al., 2007). Peptide spectrum matches (PSM) were required to be fully tryptic with any number of missed cleavages; a static modification of 57.0214 Da on cysteine (carbamidomethylated) and a dynamic modification of 15.9949 Da on methionine (oxidized) residues. PSMs were filtered using IDPicker v.3.0 (Ma et al., 2009) with an experiment-wide false-discovery rate controlled at < 1% at the peptide-level. Peptide intensities were assessed by chromatographic area-under-the-curve and unique peptide intensities summed to estimate protein-level abundance. Protein abundance distributions were then normalized across samples and missing values imputed to simulate the MS instrument’s limit of detection. Significant differences in protein abundance were assessed by pairwise T-test.

### Metabolite measurements

Strains were grown to saturation in 5 mL cultures of M9 + 2 g/L glucose + 1 mM IPTG. They were then diluted into 250 mL of M9 + 0.5 g/L glucose + 1 g/L coumarate + 1 mM IPTG and grown for a further 8 hours. The cells were separated by centrifugation, washed twice with water, and frozen in LN_2_ for later analysis.

Frozen cell pellets were weighed into centrifuge tubes containing 5 mL of 80% ethanol, and 75 μL sorbitol (1 mg/mL) added as an internal standard. Samples were sonicated for 3 min (30 s on, 30 s off with an amplitude of 30%) while being kept cold in a cooling rack that had been chilled with liquid nitrogen. Samples were then centrifuged at 4500 rpm for 20 min, the supernatant decanted and a 1 mL aliquot was dried under a nitrogen stream, dissolved in 0.5 mL acetonitrile, and silylated to generate trimethylsilyl derivatives (Tschaplinski et al., 2012). After 2 days, 1 μL aliquots were injected into an Agilent 5975C inert XL gas chromatograph-mass spectrometer (GC-MS). The standard quadrupole GC-MS was operated in electron impact (70 eV) ionization mode, targeting 2.5 full-spectrum (50-650 Da) scans per second (Tschaplinski et al., 2012).

Metabolite peaks were extracted using a key selected ion, characteristic m/z fragment to minimize integration of co-eluting metabolites. The extracted peaks of known metabolites were scaled back to the total ion current (TIC) using scaling factors previously calculated. Peaks were quantified by area integration and normalized to the quantity of internal standard recovered, amount of sample extracted, derivatized, and injected. A large user-created database and the Wiley Registry 10th Edition/NIST 2014 Mass Spectral Library was used to identify the metabolites of interest to be quantified. Unidentified metabolites were represented by their retention time and key m/z ratios.

### IMPDH expression and purification

NAD^+^ was purchased from Roche, IMP and EDTA were purchased from Fisher, MOPS was purchased from Sigma, DTT and IPTG were purchased from GoldBio. EcIMPDH/WT and AbIMPDH were purified as previously described (Makowska-Grzyska et al., 2015). pJM303, expressing EcIMPDH/D243G was transformed into BL21(*ΔguaB*) cells that lack endogenous EcIMPDH (MacPherson et al., 2010). An overnight culture (5 mL) was diluted into 1 L of fresh LB broth containing 100 μg/mL ampicillin and grown at 37 °C. Once the culture reached an OD600 of 0.6-0.8, IPTG was added to a final concentration of 0.25 mM to induce expression of IMPDH. After 13 h at 30 °C, the cells were collected by centrifugation. All the operations below were performed at 4 °C. The pellet was resuspended in 50 mL phosphate buffer (pH = 8.0) containing 1 mM dithiothreitol (DTT) and sonicated. The debris was removed by centrifugation at 10,000 g at 4 °C for 1 h.

The enzyme in the supernatant was purified by nickel affinity chromatography. The Ni-NTA resin equilibrated with water and phosphate buffer (pH = 8.0). Lysate was loaded onto the 10 mL column with 5 mL resin and washed with 50 mL phosphate buffer (pH = 8.0) then 50 mL phosphate buffer containing 25 mM imidazole. Enzyme was eluted in 25 ml phosphate buffer with 250 mM imidazole. The fractions with IMPDH activity were identified by enzyme activity assays, combined and dialyzed in 25 mM HEPES (pH = 8.0), 1mM DTT and 1 mM EDTA. Protein concentration was determined by Bio-Rad Bradford assay using IgG as a standard. The assay over-estimates the concentration of IMPDH by a factor of 2.6, and protein concentration was adjusted accordingly (Wang et al., 1996).

### IMPDH enzyme assays

The IMPDH reaction was monitored by measuring the rate of NADH production on a Shimadzu UV-1800 Spectrometer at λ = 340 nm. MOPS buffer (pH = 7.0) was used to reduce the background absorbance of 4HB. The assay buffer was composed by 20 mM MOPS (pH =7.0), 100 mM KCl, 1 mM EDTA and 1 mM DTT. The final volume of each cuvette was 1 mL.

Kinetic parameters with respect to NAD+ were determined by measuring the initial velocity for varying concentrations of NAD+ at a fixed saturating concentration of IMP (1.2 mM) and 50 nM of enzyme. Kinetic parameters with respect to IMP were determined by measuring the initial velocity for varying concentrations of IMP at a fixed saturating concentration of NAD+ (2.5 mM) and 50 nM of enzyme. Initial velocities were plotted against substrate concentrations and the data were fit using SigmaPlot. The values of *K*_i,app_ were determined by measuring the initial velocities for the reaction of IMPDH (20 nM) in the presence of varied concentrations (0-300 μM) of 4-hydroxybenzaldehyde at 12 μM of IMP and 500 μM of NAD+. The inhibition by 4-hydroxybenzaldehyde under physiological concentrations of IMP (270 μM) and NAD+ (2500 μM) were determined as well (Bennett et al., 2009; Park et al., 2016).

### Comparative modeling

HHpred (Zimmermann et al., 2017) was used to search the Protein Databank for suitable structural templates to model IMPDH from *E. coli* (accession number P0ADG7). Five templates were chosen on the basis of their similarity to the query sequence, inclusion of cofactors and substrates, or both (Table S6). Whereas all templates include the catalytic (β/α)_8_ domain, only 1ZFJ includes the cystathionine beta synthase (CBS) domain. Only 4X3Z includes the NAD+ cofactor, but it includes XMP instead of IMP. Thus, these two templates were assigned a higher weight during comparative modeling. The query and template sequences were aligned with MAFFT (L-INS-i) (Katoh and Standley, 2013) (Figure S5), and the wild-type IMPDH sequence was threaded onto each template structure.

We then used RosettaCM (Song et al., 2013) to generate comparative models of IMPDH. Fragment files were obtained with the Robetta web server (Gront et al., 2011; Kim et al., 2004). We used an iterative approach in which we first generated 1,000 models with all five templates. We then selected the top model along with the 4X3Z and 1ZFJ templates and generated an additional 1,000 models. Three iterations were carried out to obtain the final model used for ligand docking.

### Ligand docking

Structure files in mol2 format for IMP (ZINC04228242), NAD^+^ (ZINC08214766), and 4-hydroxybenzaldehyde (ZINC00156709) were obtained from http://zinc.docking.org (Irwin et al., 2012) and then converted to Rosetta format. RosettaLigand (Meiler and Baker, 2006) was then used to dock the inhibitor 4-hydroxybenzaldehyde into the active site of IMPDH following a previously described protocol (Combs et al., 2013). Top binding poses were ranked on the basis of their ‘interface_delta_X’ score in Rosetta energy units. Additional details of the comparative modeling and ligand docking are provided in the Supporting Information.

## Acknowledgments

Genome resequencing and analysis was performed by Christa Pennacchio, Natasha Brown, Anna Lipzen, and Wendy Schackwitz at the Joint Genome Institute. DNA synthesis was performed by Jan-Fang Cheng, Samuel Deutsch, and Miranda Harmon-Smith at the Joint Genome Institute. The work conducted by the U.S. Department of Energy Joint Genome Institute, a DOE Office of Science User Facility, is supported by the Office of Science of the U.S. Department of Energy under Contract No. DE-AC02-05CH11231. This work was supported by the BioEnergy Science Center and The Center for Bioenergy Innovation, both U.S. Department of Energy Research Centers supported by the Office of Biological and Environmental Research in the DOE Office of Science; the National Institutes of Health (GM054403 to LH); and the ORNL Laboratory Directed Research and Development program (#8949 to JMP). SJC was supported by NIH/NIGMS-IMSD Grant No. R25GM086761. This material is based upon work supported by the National Science Foundation Graduate Research Fellowship under Grant No. (2017219379). Oak Ridge National Laboratory (ORNL) is managed by UT-Battelle, LLC, for the DOE under Contract No. DE-AC05-00OR22725.

---

**Table S1:**
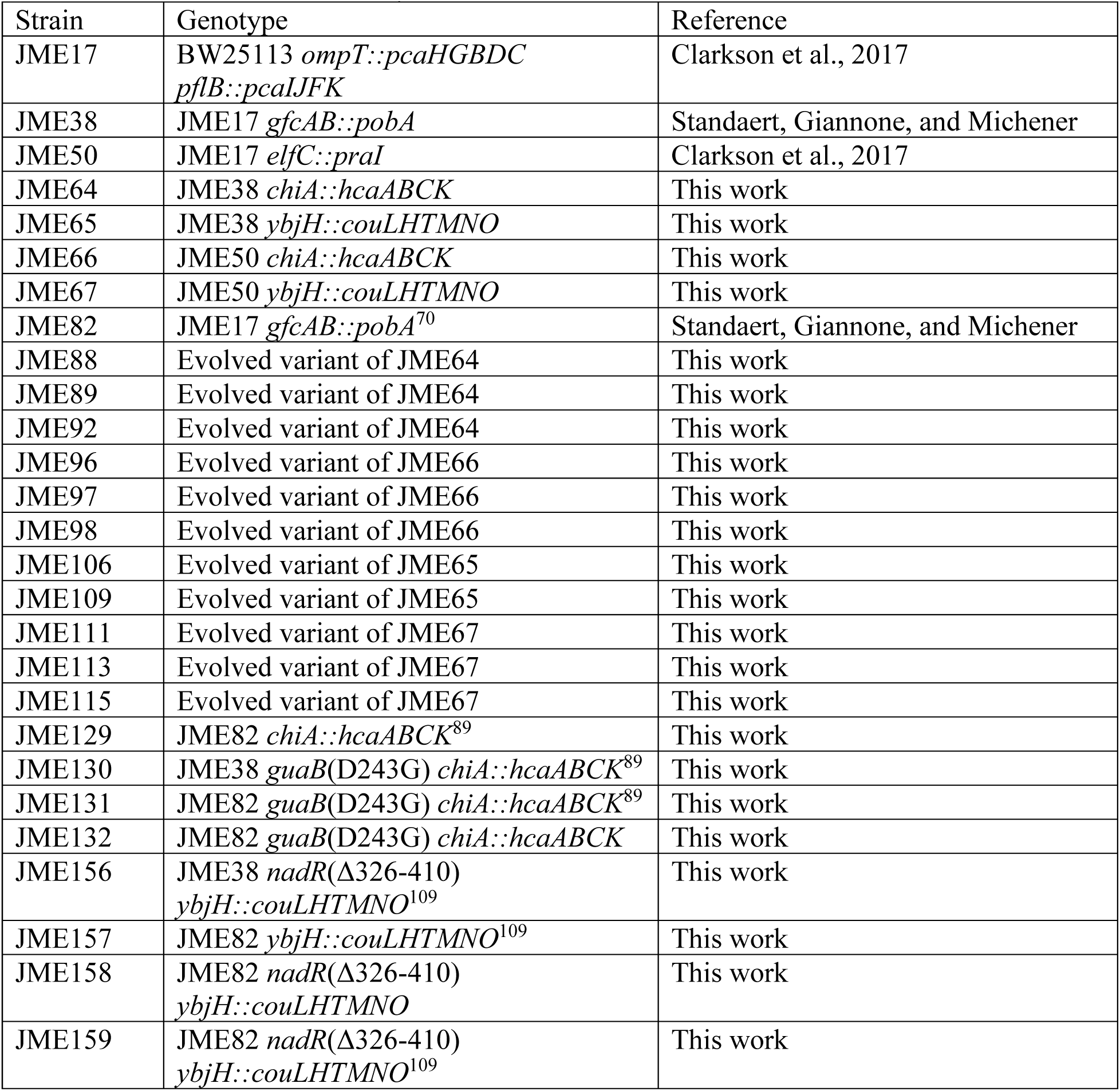
Strains used in this study

**Table S2:** Plasmids used in this study

| Plasmid | Description | Reference |
| --- | --- | --- |
| pCas | Expresses Cas9 and λ-RED | Jiang et al., 2015 |
| pTarget | Expresses sgRNA | Jiang et al., 2015 |
| pJM157 | Expresses *elfC* sgRNA | Clarkson et al., 2017 |
| pJM168 | Expresses *gfcAB* sgRNA | Standaert, Giannone, and Michener |
| pJM187 | Expresses *yjbH* sgRNA | This work |
| pJM205 | Expresses *chiA* sgRNA | This work |
| pJM262 | Expresses *guaB* sgRNA | This work |
| pJM284 | Expresses *nadR* sgRNA | This work |
| pJM303 | pMCSG7 *guaB* | This work |

**Table S3:**
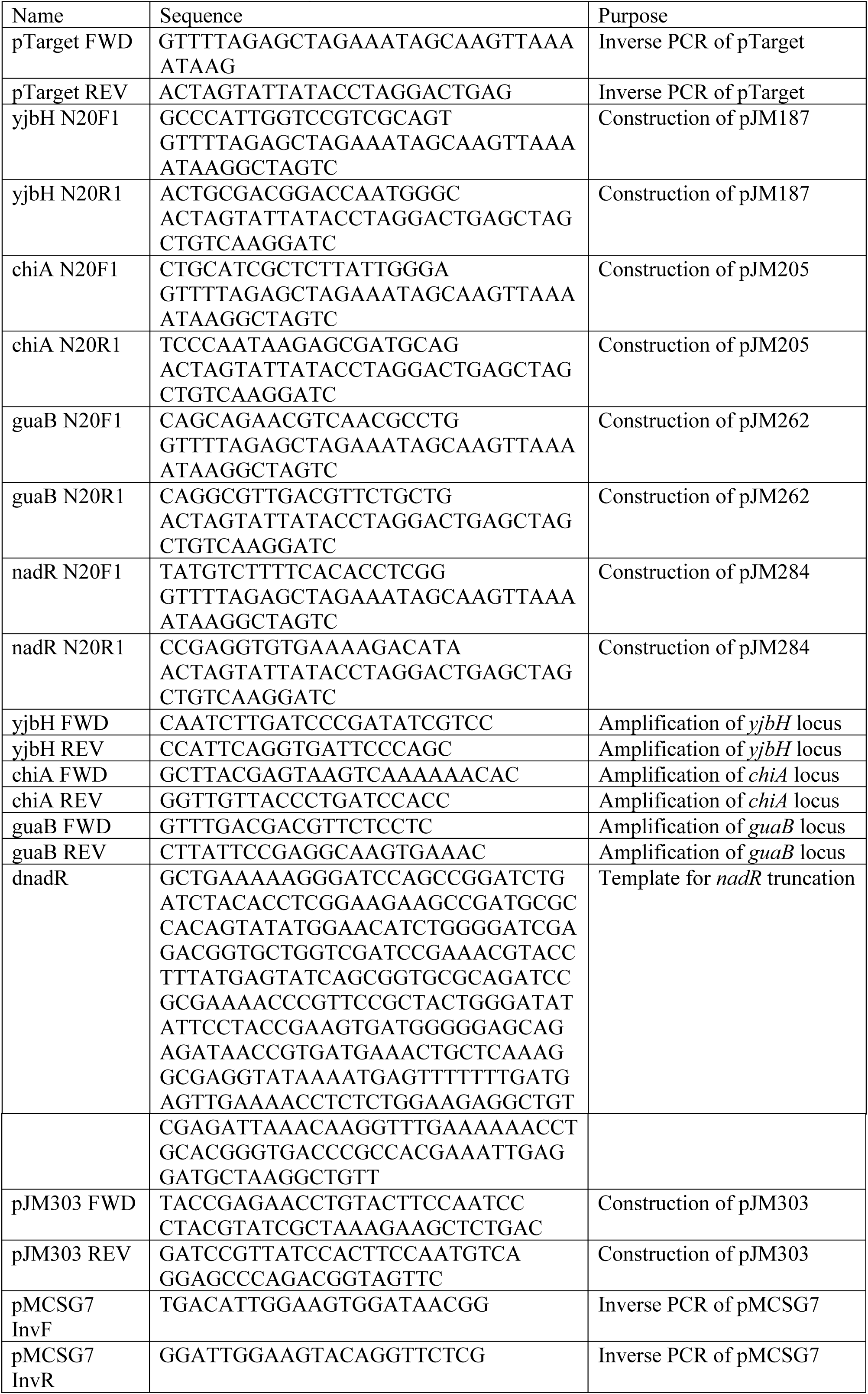
Primers used in this study

**Table S4:**
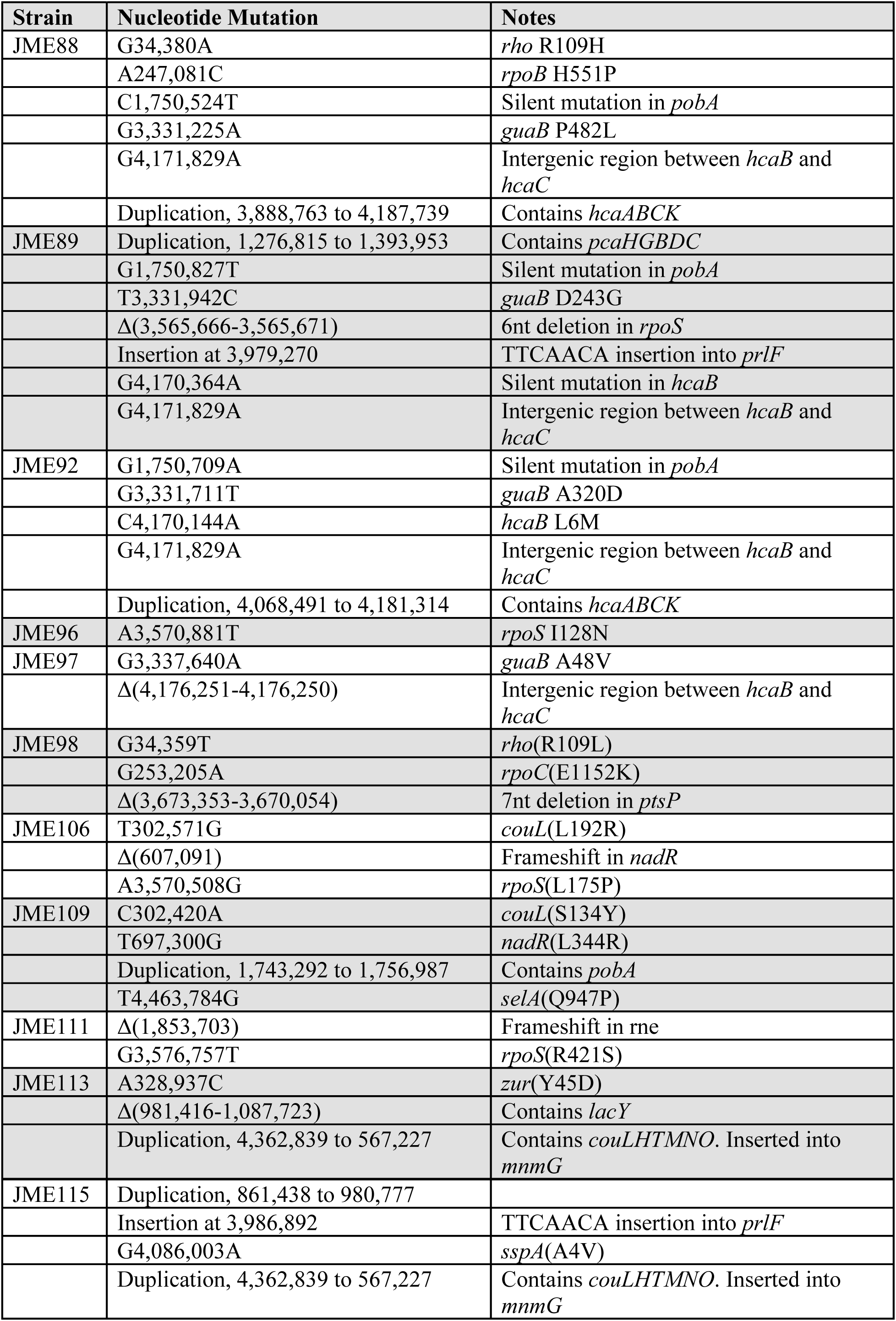
Mutations identified. All nucleotide references are relative to the appropriate parental strain.

**Table S5:**
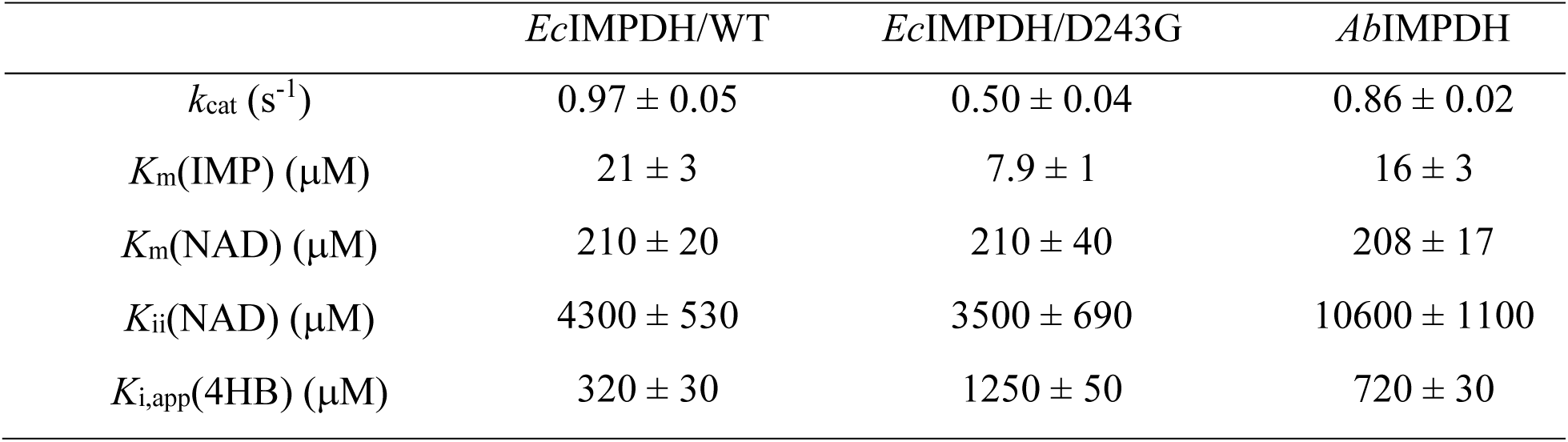
Kinetic Parameters for *Ec*IMPDH/WT, *Ec*IMPDH/D243G and *Ab*IMPDH. The values are the average and range of two independent experiments.

|  | *Ec*IMPDH/WT | *Ec*IMPDH/D243G | *Ab*IMPDH |
| --- | --- | --- | --- |
| $k_{\text{cat}}$ (s<sup>-1</sup>) | 0.97 ± 0.05 | 0.50 ± 0.04 | 0.86 ± 0.02 |
| $K_{\text{m}}$(IMP) (μM) | 21 ± 3 | 7.9 ± 1 | 16 ± 3 |
| $K_{\text{m}}$(NAD) (μM) | 210 ± 20 | 210 ± 40 | 208 ± 17 |
| $K_{\text{ii}}$(NAD) (μM) | 4300 ± 530 | 3500 ± 690 | 10600 ± 1100 |
| $K_{\text{i,app}}$(4HB) (μM) | 320 ± 30 | 1250 ± 50 | 720 ± 30 |

**Table S6:** Crystallographic templates used to model IMPDH from *E. coli*

| PDB entry | Organism | Resolution (Å) | E value | % identity | Cofactors/ligands |
| --- | --- | --- | --- | --- | --- |
| 4X3Z | *Vibrio cholerae* | 1.62 | 3.3e-34 | 86 | NAD<sup>+</sup>, XMP |
| 1ZFJ | *Streptococcus pyogenes* | 1.90 | 9.8e-55 | 56 | IMP |
| 5AHN | *Pseudomonas aeruginosa* | 1.65 | 4.8e-59 | 66 | IMP |
| 2CU0 | *Pyrococcus horikoshii* | 2.10 | 2.9e-47 | 49 | XMP |
| 1VRD | *Thermotoga maritima* | 2.18 | 8.3e-47 | 56 | n/a |

**Table S7:** Binding energies^a^ for the top five poses obtained from docking 4-hydroxybenzaldehyde to the apoenzyme, IMP-bound, and IMP/NAD^+^-bound states of IMPDH.

| *interface_delta* (Rosetta energy units) |  |  |
| --- | --- | --- |
| apoenzyme | IMP-bound | IMP/NAD<sup>+</sup>-bound |
| -11.5 | -12.4 | -10.3 |
| -11.0 | -12.2 | -9.4 |
| -10.6 | -12.1 | -9.3 |
| -10.6 | -12.1 | -8.9 |
| -10.5 | -11.4 | -8.7 |

<sup>a</sup> *interface\_delta*

**Figure S1:**
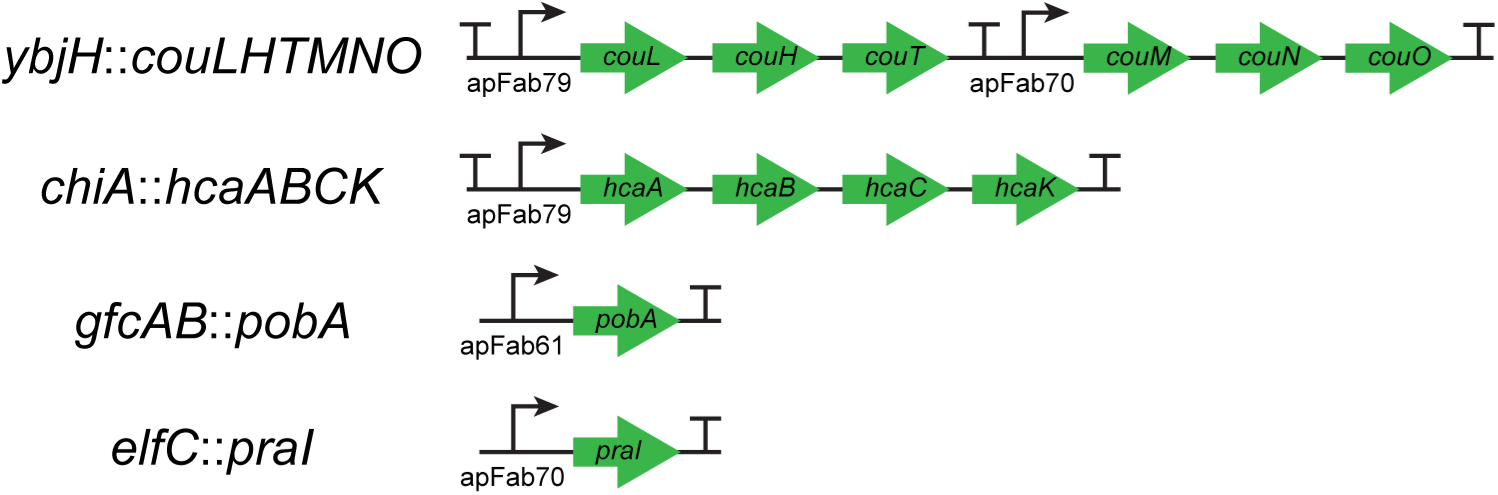
Construct designs. Sequences were synthesized *de novo* and inserted into the indicated chromosomal locus.

**Figure S2:**
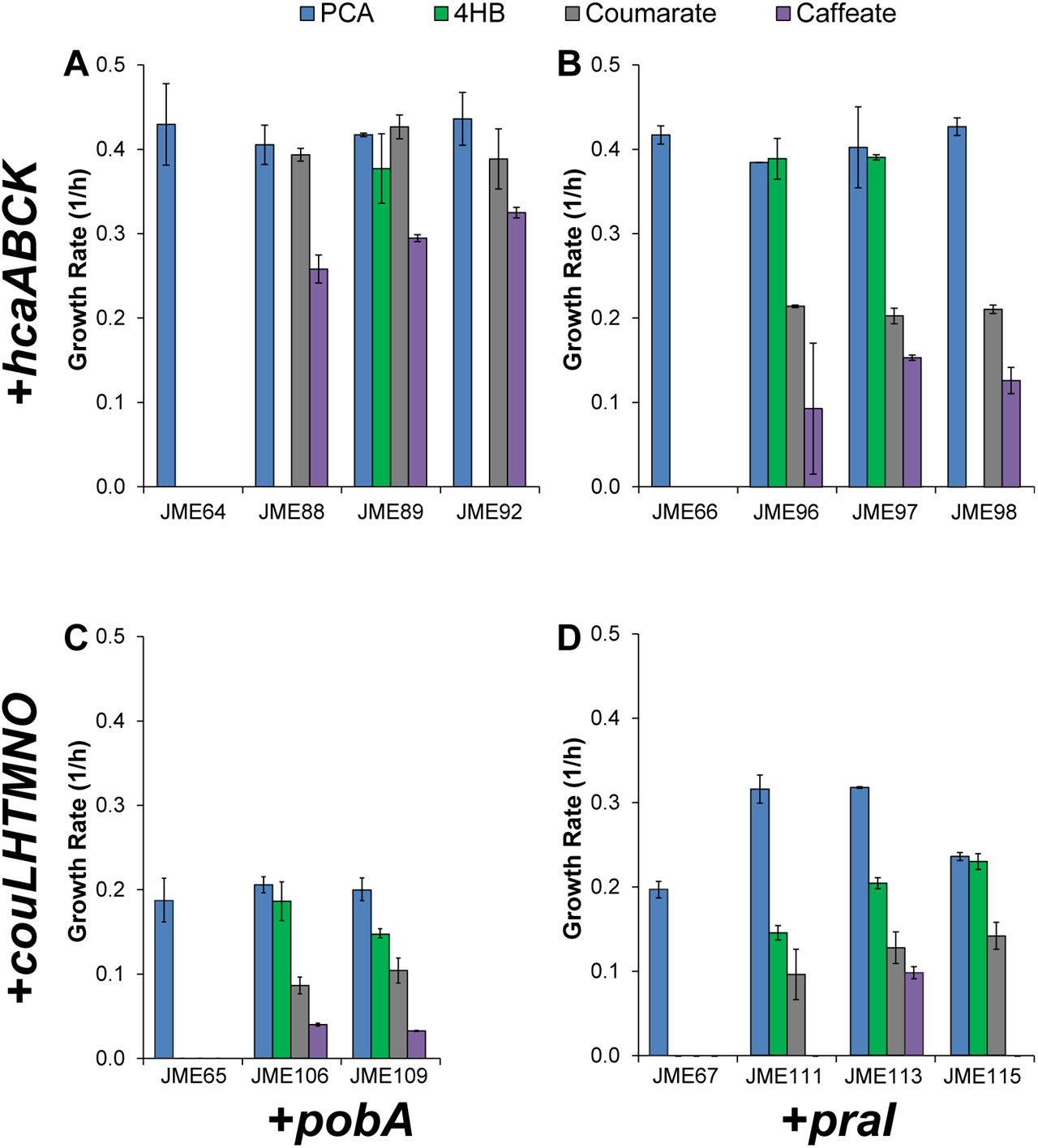
A combination of engineering and evolution are necessary for growth with coumarate. Engineered strains (JME64, JME65, JME66, and JME67) and their evolved derivatives were grown in minimal media containing 1 g/L of the indicated substrate. Each panel represents a different combination of phenylpropanoid pathway (A and B: *hca*, C and D: *cou*) and 4-HB monooxygenase (A and C: *pobA*, B and D: *praI*). Error bars show one standard deviation, calculated from three biological replicates.

**Figure S3:**
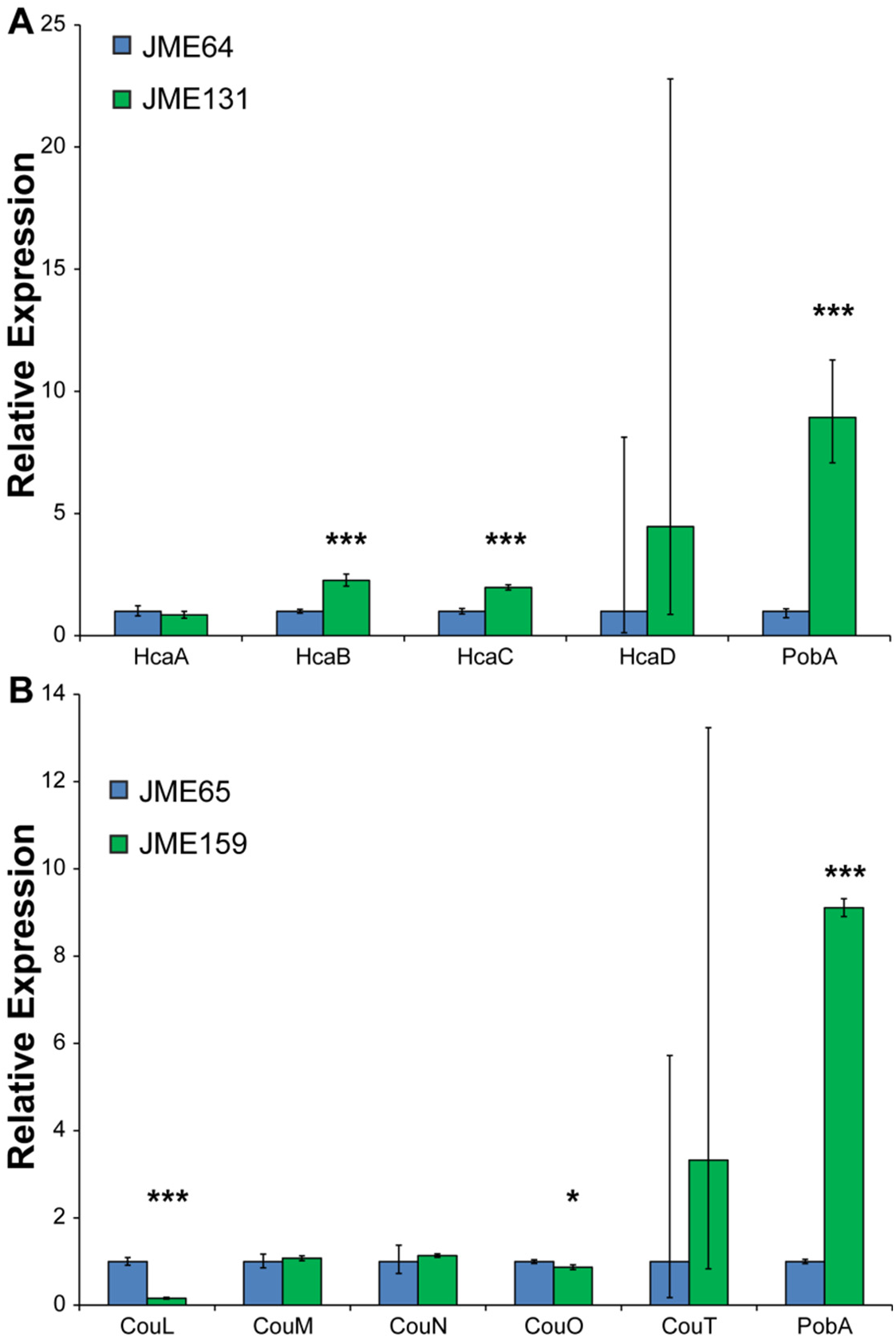
Pathway mutations affect enzyme expression. (A) In strains with the *hca* pathway, an intergenic mutation between *hcaB* and *hcaC* increases expression of both enzymes, while a silent mutation to *pobA* also increases expression of that enzyme. Strain JME64 has the wild-type allele for both constructs, while JME131 has both mutations. Error bars show one standard deviation, calculated from three biological replicates. ***: p<0.001.

**Figure S4:**
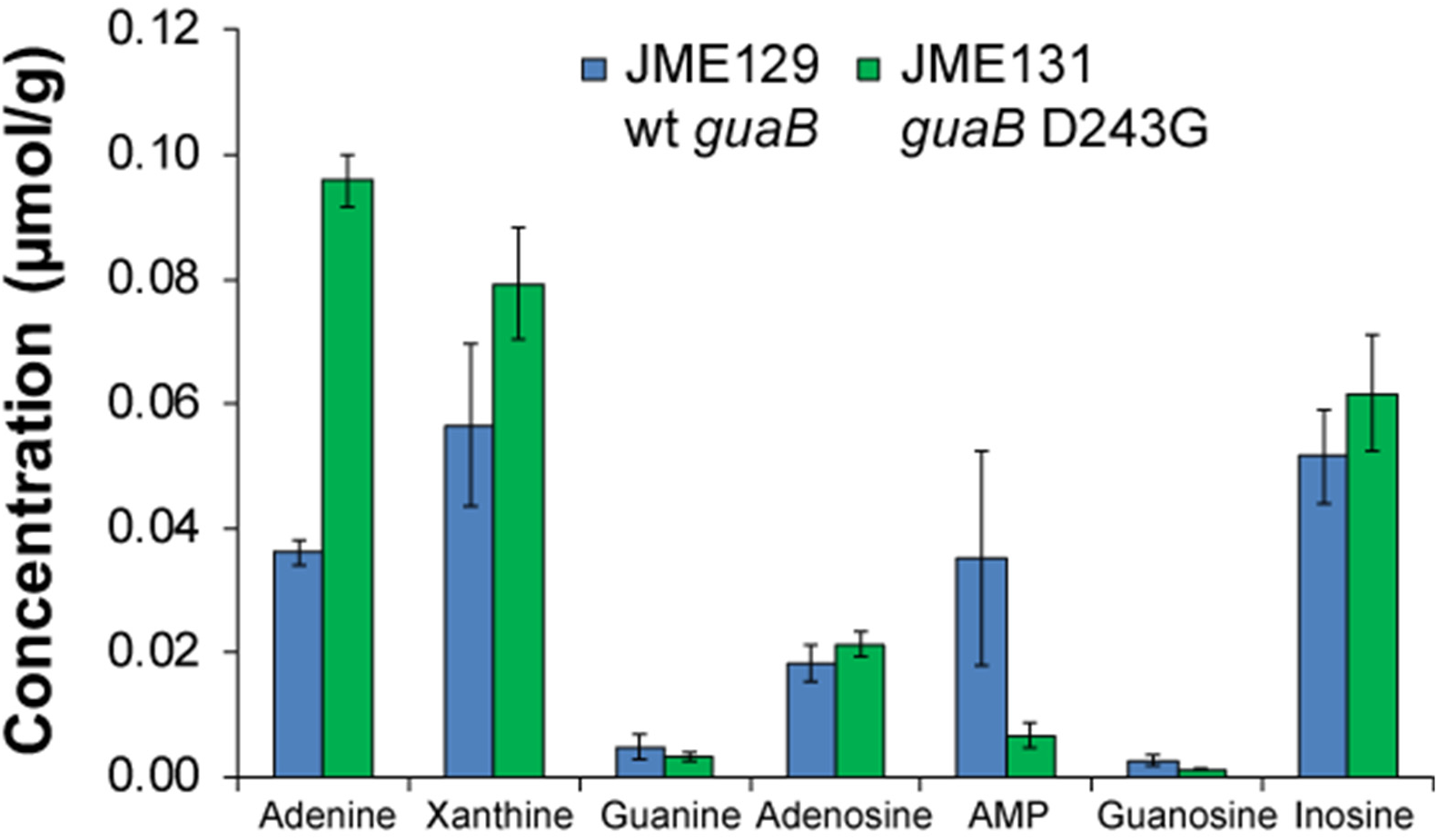
Metabolomics of purine nucleobases and derivatives. Concentrations of GMP, XMP, and IMP were below the limit of detection. Concentrations are reported in μmol/g fresh weight sorbitol equivalent. Error bars show one standard deviation, calculated from four biological replicates.

**Figure S5:**
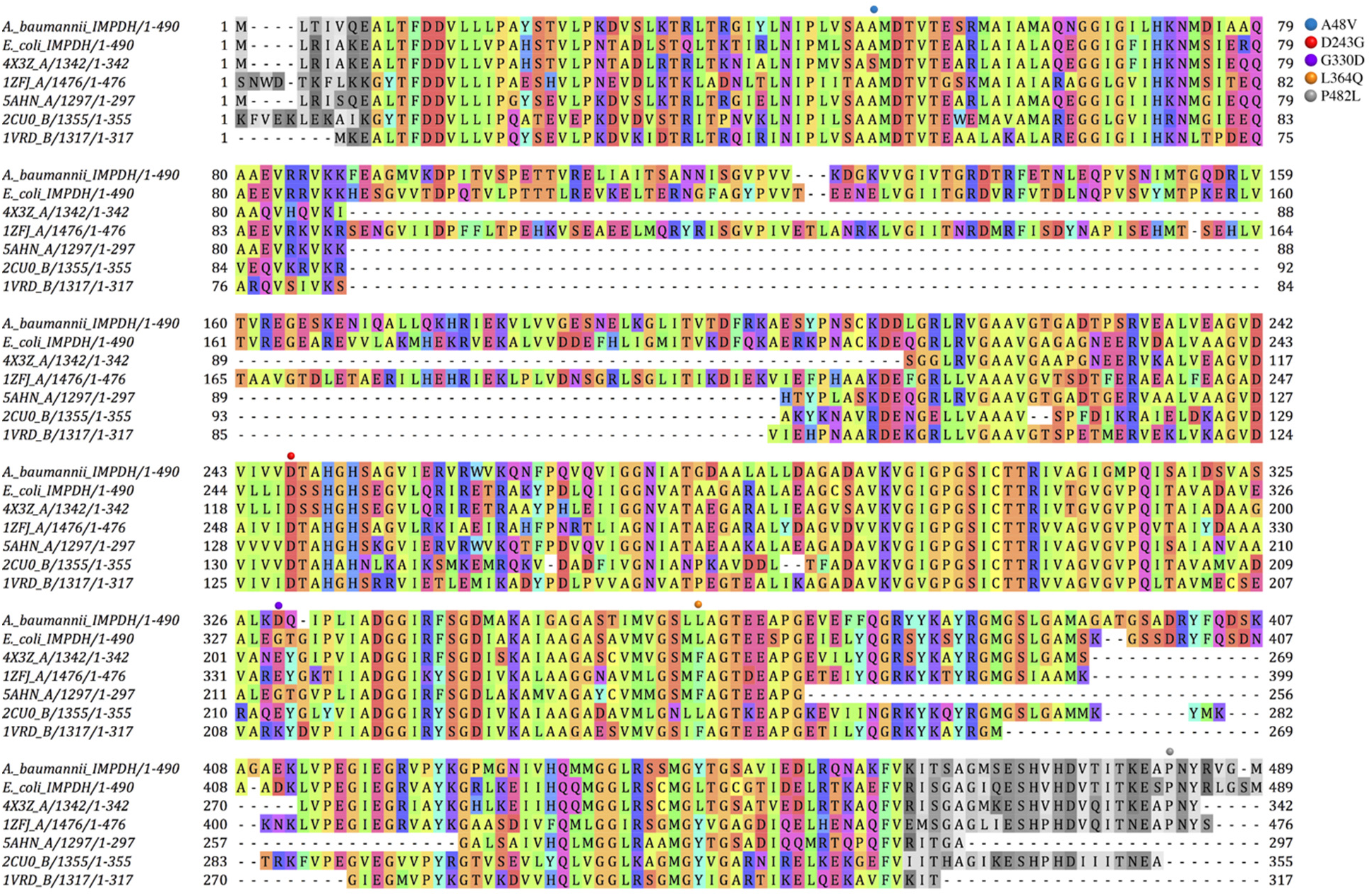
MAFFT (L-INS-i) multiple sequence alignment of IMPDH from *A. baumannii*, *E. coli*, and multiple template sequences used for structural modeling of the *E. coli* IMPDH. Selected residues (gray) were trimmed at the N- and C-termini and were not included in the models.

**Figure S6:**
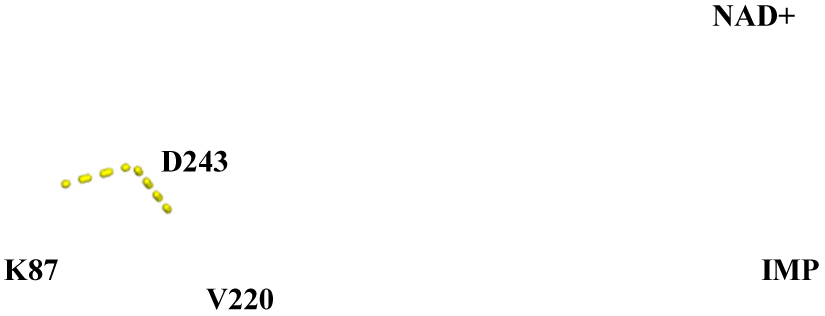
Local environment of D243 in the model of *E. coli* wild-type IMPDH. The carboxylate side chain of D243 forms hydrogen bonds with the side chain of K87 and the backbone of V220. K87 is located at the C-terminal end of a long α helix, and V220 is at the beginning of a β strand (shown in maroon). Carbons of K87, V220 and D243 are shown in gray, and carbons of IMP and NAD^+^ are shown in yellow.

## Example options file for comparative modeling

~~~
-in:file:fasta input/t000_.fasta
-parser:protocol input/rosetta_cm.xml
-relax:minimize_bond_angles
-relax:minimize_bond_lengths
-default_max_cycles 200
-relax:min_type lbfgs_armijo_nonmonotone
-use_bicubic_interpolation
-relax:jump_move true
-score:weights input/ref2015_cart.wts
-hybridize:stage1_probability 1.0
-mute all
-nstruct 1000
~~~

## Example XML file for comparative modeling

~~~
<ROSETTASCRIPTS>
        <TASKOPERATIONS>
        </TASKOPERATIONS>
        <SCOREFXNS>
               <ScoreFunction name=“stage1” weights=“input/stage1.wts” symmetric=“1”>
                    <Reweight scoretype=“atom_pair_constraint” weight=“1”/>
               </ScoreFunction>
               <ScoreFunction name=“stage2” weights=“input/stage2.wts” symmetric=“1”>
                   <Reweight scoretype=“atom_pair_constraint” weight=“0.5” />
               </ScoreFunction>
               <ScoreFunction name=“fullatom” weights=“input/stage3.wts” symmetric=“1”>
                   <Reweight scoretype=“atom_pair_constraint” weight=“0.5” />
              </ScoreFunction>
     </SCOREFXNS>
<FILTERS>
</FILTERS>
<MOVERS>
        <Hybridize name=“hybridize” stage1_scorefxn=“stage1” stage2_scorefxn=“stage2” fa_scorefxn=“fullatom” batch=“1” stage1_increase_cycles=“1.0” stage2_increase_cycles=“1.0” linmin_only=“1”>
              <Template pdb=“input/impdh_on_4X3Z_A.pdb” cst_file=“AUTO” weight=“1.000” symmdef=“input/tetramer_4x3z.symm”/>
              <Template pdb=“input/impdh_on_1ZFJ_A.pdb” cst_file=“AUTO” weight=“1.000” symmdef=“input/tetramer_1zfj.symm”/>
              <Template pdb=“input/try7_cm1_S_0278_A.pdb” cst_file=“AUTO” weight=“1.000” symmdef=“input/tetramer_S_0278.symm”/>
              <Fragments three_mers=“input/aat000_03_05.200_v1_3.txt.gz” nine_mers=“input/aat000_09_05.200_v1_3.txt.gz”/>
        </Hybridize>
</MOVERS>
<APPLY_TO_POSE></APPLY_TO_POSE>
<PROTOCOLS>
          <Add mover=“hybridize”>
          </Add>
          </PROTOCOLS>
</ROSETTASCRIPTS>
~~~

## Example options file for ligand docking

~~~
-in:file:s ‘S_0308_aligned_with_4X3Z_A_1141_zinc.pdb’
-in:file:extra_res_fa IMP.params NAD.params zinc_156709.params
-packing:ex1
-packing:ex2
-packing:no_optH false
-packing:flip_HNQ true
-packing:ignore_ligand_chi true
-parser:protocol dock.xml
-mistakes:restore_pre_talaris_2013_behavior true
-overwrite
-nstruct 1000
-mute all
~~~

## Example XML file for ligand docking

~~~
<ROSETTASCRIPTS>
        <SCOREFXNS>
                <ScoreFunction name=“ligand_soft_rep” weights=“ligand_soft_rep”>
                </ScoreFunction>
                <ScoreFunction name=“hard_rep” weights=“ligand”>
                </ScoreFunction>
        </SCOREFXNS>
        <LIGAND_AREAS>
                <LigandArea name=“inhibitor_dock_sc” chain=“X” cutoff=“6.0” add_nbr_radius=“true” all_atom_mode=“false”/>
                <LigandArea name=“inhibitor_final_sc” chain=“X” cutoff=“6.0” add_nbr_radius=“true” all_atom_mode=“false”/>
                <LigandArea name=“inhibitor_final_bb” chain=“X” cutoff=“7.0” add_nbr_radius=“false” all_atom_mode=“true” Calpha_restraints=“0.3”/>
        </LIGAND_AREAS>
        <INTERFACE_BUILDERS>
                <InterfaceBuilder name=“side_chain_for_docking” ligand_areas=“inhibitor_dock_sc”/>
                <InterfaceBuilder name=“side_chain_for_final” ligand_areas=“inhibitor_final_sc”/>
                <InterfaceBuilder name=“backbone” ligand_areas=“inhibitor_final_bb” extension_window=“3”/>
        </INTERFACE_BUILDERS>
        <MOVEMAP_BUILDERS>
                <MoveMapBuilder name=“docking” sc_interface=“side_chain_for_docking” minimize_water=“false”/>
                <MoveMapBuilder name=“final” sc_interface=“side_chain_for_final” bb_interface=“backbone” minimize_water=“false”/>
        </MOVEMAP_BUILDERS>
        <SCORINGGRIDS ligand_chain=“X” width=“15”>
                <ClassicGrid grid_name=“classic” weight=“1.0”/>
        </SCORINGGRIDS>
        <MOVERS>
                <StartFrom name=“start_from_X” chain=“X”>
                        <Coordinates x=“-25.1” y=“21.2” z=“20.4”/>
               </StartFrom>
               <Transform name=“transform” chain=“X” box_size=“20.0” move_distance=“0.2” angle=“20” cycles=“500” repeats=“1” temperature=“5”/>
               <HighResDocker name=“high_res_docker” cycles=“6” repack_every_Nth=“3” scorefxn=“ligand_soft_rep” movemap_builder=“docking”/>
               <FinalMinimizer name=“final” scorefxn=“hard_rep” movemap_builder=“final”/>
               <InterfaceScoreCalculator name=“add_scores” chains=“X” scorefxn=“hard_rep” />
       </MOVERS>
       <PROTOCOLS>
               <Add mover_name=“start_from_X”/>
               <Add mover_name=“transform”/>
               <Add mover_name=“high_res_docker”/>
               <Add mover_name=“final”/>
               <Add mover_name=“add_scores”/>
       </PROTOCOLS>
</ROSETTASCRIPTS>
~~~

## References

Bennett, B.D., Kimball, E.H., Gao, M., Osterhout, R., Van Dien, S.J., Rabinowitz, J.D., 2009. Absolute metabolite concentrations and implied enzyme active site occupancy in Escherichia coli. Nat. Chem. Biol. 5, 593–599. doi:10.1038/nchembio.186

Bomble, Y.J., Lin, C.-Y., Amore, A., Wei, H., Holwerda, E.K., Ciesielski, P.N., Donohoe, B.S., Decker, S.R., Lynd, L.R., Himmel, M.E., 2017. Lignocellulose deconstruction in the biosphere. Curr. Opin. Chem. Biol. 41, 61–70. doi:10.1016/J.CBPA.2017.10.013

Bugg, T.D.H., Ahmad, M., Hardiman, E.M., Rahmanpour, R., 2011. Pathways for degradation of lignin in bacteria and fungi. Nat. Prod. Rep. 28, 1883. doi:10.1039/c1np00042j

Chen, Y.-J., Liu, P., Nielsen, A.A.K., Brophy, J.A.N., Clancy, K., Peterson, T., Voigt, C.A., 2013. Characterization of 582 natural and synthetic terminators and quantification of their design constraints. Nat. Methods. 10, 659–664. doi:10.1038/nmeth.2515

Clark, I.C., Melnyk, R.A., Youngblut, M.D., Carlson, H.K., Iavarone, A.T., Coates, J.D., 2015. Synthetic and evolutionary construction of a chlorate-reducing *Shewanella oneidensis* MR-1. MBio 6, e00282–15. doi:10.1128/mBio.00282-15

Clarkson, S.M., Hamilton-Brehm, S.D., Giannone, R.J., Engle, N.L., Tschaplinski, T.J., Hettich, R.L., Elkins, J.G., 2014. A comparative multidimensional LC-MS proteomic analysis reveals mechanisms for furan aldehyde detoxification in Thermoanaerobacter pseudethanolicus 39E. Biotechnol. Biofuels. 7, 165. doi:10.1186/s13068-014-0165-z

Clarkson, S.M., Kridelbaugh, D.M., Elkins, J.G., Guss, A.M., Michener, J., 2017. Construction and optimization of a heterologous pathway for protocatechuate catabolism in Escherichia coli enables rapid bioconversion of model lignin monomers. Appl. Environ. Microbiol.. 83, e01313–17. doi:10.1128/AEM.01313-17

Combs, S.A., DeLuca, S.L., DeLuca, S.H., Lemmon, G.H., Nannemann, D.P., Nguyen, E.D., Willis, J.R., Sheehan, J.H., Meiler, J., 2013. Small-molecule ligand docking into comparative models with Rosetta. Nat. Protoc. 8, 1277–1298. doi:10.1038/nprot.2013.074

Delaney, N.F., Kaczmarek, M.E., Ward, L.M., Swanson, P.K., Lee, M.-C., Marx, C.J., 2013. Development of an optimized medium, strain and high-throughput culturing methods for Methylobacterium extorquens. PLoS One. 8, e62957. doi:10.1371/journal.pone.0062957

Espah Borujeni, A., Channarasappa, A.S., Salis, H.M., 2014. Translation rate is controlled by coupled trade-offs between site accessibility, selective RNA unfolding and sliding at upstream standby sites. Nucleic Acids Res. 42, 2646–59. doi:10.1093/nar/gkt1139

Gront, D., Kulp, D.W., Vernon, R.M., Strauss, C.E.M., Baker, D., 2011. Generalized Fragment Picking in Rosetta: Design, Protocols and Applications. PLoS One. 6, e23294. doi:10.1371/journal.pone.0023294

Hedstrom, L., 2009. IMP dehydrogenase: structure, mechanism, and inhibition. Chem. Rev. 109, 2903–28. doi:10.1021/cr900021w

Irwin, J.J., Sterling, T., Mysinger, M.M., Bolstad, E.S., Coleman, R.G., 2012. ZINC: A Free Tool to Discover Chemistry for Biology. J. Chem. Inf. Model. 52, 1757–1768. doi:10.1021/ci3001277

Jiang, Y., Chen, B., Duan, C., Sun, B., Yang, J., Yang, S., 2015. Multigene editing in the Escherichia coli genome via the CRISPR-Cas9 system. Appl. Environ. Microbiol. 81, 2506–2514. doi:10.1128/AEM.04023-14

Katoh, K., Standley, D.M., 2013. MAFFT Multiple Sequence Alignment Software Version 7: Improvements in Performance and Usability. Mol. Biol. Evol. 30, 772–780. doi:10.1093/molbev/mst010

Kim, D.E., Chivian, D., Baker, D., 2004. Protein structure prediction and analysis using the Robetta server. Nucleic Acids Res. 32, W526–W531. doi:10.1093/nar/gkh468

Kim, J., Copley, S.D., 2012. Inhibitory cross-talk upon introduction of a new metabolic pathway into an existing metabolic network. Proc. Natl. Acad. Sci. 109, E2856–E2864. doi:10.1073/pnas.1208509109

Kizer, L., Pitera, D.J., Pfleger, B.F., Keasling, J.D., 2008. Application of Functional Genomics to Pathway Optimization for Increased Isoprenoid Production. Appl. Environ. Microbiol. 74, 3229–3241. doi:10.1128/AEM.02750-07

Kosuri, S., Goodman, D.B., Cambray, G., Mutalik, V.K., Gao, Y., Arkin, A.P., Endy, D., Church, G.M., 2013. Composability of regulatory sequences controlling transcription and translation in *Escherichia coli*. Proc. Natl. Acad. Sci. U. S. A. 110, 14024–9. doi:10.1073/pnas.1301301110

Kurnasov, O. V, Polanuyer, B.M., Ananta, S., Sloutsky, R., Tam, A., Gerdes, S.Y., Osterman, A.L., 2002. Ribosylnicotinamide kinase domain of NadR protein: identification and implications in NAD biosynthesis. J. Bacteriol. 184, 6906–17. doi:10.1128/JB.184.24.6906-6917.2002

Linger, J.G., Vardon, D.R., Guarnieri, M.T., Karp, E.M., Hunsinger, G.B., Franden, M.A., Johnson, C.W., Chupka, G., Strathmann, T.J., Pienkos, P.T., Beckham, G.T., 2014. Lignin valorization through integrated biological funneling and chemical catalysis. Proc. Natl. Acad. Sci. U. S. A. 111, 12013–8. doi:10.1073/pnas.1410657111

Ma, Z.-Q., Dasari, S., Chambers, M.C., Litton, M.D., Sobecki, S.M., Zimmerman, L.J., Halvey, P.J., Schilling, B., Drake, P.M., Gibson, B.W., Tabb, D.L., 2009. IDPicker 2.0: Improved protein assembly with high discrimination peptide identification filtering. J. Proteome Res. 8, 3872–3881. doi:10.1021/pr900360j

MacPherson, I.S., Kirubakaran, S., Gorla, S.K., Riera, T. V., D’Aquino, J.A., Zhang, M., Cuny, G.D., Hedstrom, L., 2010. The Structural Basis of Cryptosporidium-Specific IMP Dehydrogenase Inhibitor Selectivity. J. Am. Chem. Soc. 132, 1230–1231. doi:10.1021/ja909947a

Makowska-Grzyska, M., Kim, Y., Maltseva, N., Osipiuk, J., Gu, M., Zhang, M., Mandapati, K., Gollapalli, D.R., Gorla, S.K., Hedstrom, L., Joachimiak, A., 2015. A Novel Cofactor-binding Mode in Bacterial IMP Dehydrogenases Explains Inhibitor Selectivity. J. Biol. Chem. 290, 5893–5911. doi:10.1074/jbc.M114.619767

Meiler, J., Baker, D., 2006. ROSETTALIGAND: Protein-small molecule docking with full side-chain flexibility. Proteins Struct. Funct. Bioinforma. 65, 538–548. doi:10.1002/prot.21086

Michener, J.K., Camargo Neves, A.A., Vuilleumier, S., Bringel, F., Marx, C.J., 2014. Effective use of a horizontally-transferred pathway for dichloromethane catabolism requires post-transfer refinement. Elife. 3. doi:10.7554/eLife.04279

Michener, J.K., Nielsen, J., Smolke, C.D., 2012. Identification and treatment of heme depletion attributed to overexpression of a lineage of evolved P450 monooxygenases. Proc. Natl. Acad. Sci. 109, 19504–19509. doi:10.1073/pnas.1212287109

Mills, T.Y., Sandoval, N.R., Gill, R.T., 2009. Cellulosic hydrolysate toxicity and tolerance mechanisms in Escherichia coli. Biotechnol. Biofuels. 2, 26. doi:10.1186/1754-6834-2-26

Mukhopadhyay, A., 2015. Tolerance engineering in bacteria for the production of advanced biofuels and chemicals. Trends Microbiol. 23, 498–508. doi:http://dx.doi.org/10.1016/j.tim.2015.04.008

Nielsen, J., Keasling, J.D., 2016. Engineering Cellular Metabolism. Cell. 164, 1185–1197. doi:10.1016/j.cell.2016.02.004

Otani, H., Lee, Y.-E., Casabon, I., Eltis, L.D., 2014. Characterization of p-hydroxycinnamate catabolism in a soil Actinobacterium. J. Bacteriol. 196, 4293–303. doi:10.1128/JB.02247-14

Pál, C., Papp, B., Lercher, M.J., 2005. Adaptive evolution of bacterial metabolic networks by horizontal gene transfer. Nat. Genet. 37, 1372–1375. doi:10.1038/ng1686

Park, J.O., Rubin, S.A., Xu, Y.-F., Amador-Noguez, D., Fan, J., Shlomi, T., Rabinowitz, J.D., 2016. Metabolite concentrations, fluxes and free energies imply efficient enzyme usage. Nat. Chem. Biol. 12, 482–489. doi:10.1038/nchembio.2077

Parke, D., Ornston, L.N., 2003. Hydroxycinnamate (hca) Catabolic Genes from Acinetobacter sp. Strain ADP1 Are Repressed by HcaR and Are Induced by Hydroxycinnamoyl-Coenzyme A Thioesters. Appl. Environ. Microbiol. 69, 5398–5409. doi:10.1128/AEM.69.9.5398-5409.2003

Pisithkul, T., Jacobson, T.B., O’Brien, T.J., Stevenson, D.M., Amador-Noguez, D., 2015. Phenolic Amides Are Potent Inhibitors of De Novo Nucleotide Biosynthesis. Appl. Environ. Microbiol. 81, 5761–72. doi:10.1128/AEM.01324-15

Porse, A., Schou, T.S., Munck, C., Ellabaan, M.M.H., Sommer, M.O.A., 2018. Biochemical mechanisms determine the functional compatibility of heterologous genes. Nat. Commun. 9, 522. doi:10.1038/s41467-018-02944-3

Ragauskas, A.J., Beckham, G.T., Biddy, M.J., Chandra, R., Chen, F., Davis, M.F., Davison, B.H., Dixon, R.A., Gilna, P., Keller, M., Langan, P., Naskar, A.K., Saddler, J.N., Tschaplinski, T.J., Tuskan, G.A., Wyman, C.E., 2014. Lignin valorization: improving lignin processing in the biorefinery. Science. 344, 1246843. doi:10.1126/science.1246843

Rodriguez, A., Salvachua, D., Katahira, R., Black, B.A., Cleveland, N.S., Reed, M.L., Smith, H., Baidoo, E.E.K., Keasling, J.D., Simmons, B.A., Beckham, G.T., Gladden, J.M., 2017. Base-catalyzed depolymerization of solid lignin-rich streams enables microbial conversion. ACS Sustain. Chem. Eng. acssuschemeng. 7b01818. doi:10.1021/acssuschemeng.7b01818

Salis, H.M., Mirsky, E.A., Voigt, C.A., 2009. Automated design of synthetic ribosome binding sites to control protein expression. Nat. Biotechnol. 27, 946–950. doi:10.1038/nbt.1568

Saxer, G., Krepps, M.D., Merkley, E.D., Ansong, C., Deatherage Kaiser, B.L., Valovska, M.-T., Ristic, N., Yeh, P.T., Prakash, V.P., Leiser, O.P., Nakhleh, L., Gibbons, H.S., Kreuzer, H.W., Shamoo, Y., 2014. Mutations in Global Regulators Lead to Metabolic Selection during Adaptation to Complex Environments. PLoS Genet.. 10, e1004872. doi:10.1371/journal.pgen.1004872

Song, Y., DiMaio, F., Wang, R.Y.-R., Kim, D., Miles, C., Brunette, T., Thompson, J., Baker, D., 2013. High-Resolution Comparative Modeling with RosettaCM. Structure. 21, 1735–1742. doi:10.1016/J.STR.2013.08.005

Standaert, R.F., Giannone, R.J., Michener, J.K., 2018. Identification of parallel and divergent optimization solutions for homologous metabolic enzymes. Metab. Eng. Commun. 6, 56–62. doi:10.1016/J.METENO.2018.04.002

Tabb, D.L., Fernando, C.G., Chambers, M.C., 2007. MyriMatch: highly accurate tandem mass spectral peptide identification by multivariate hypergeometric analysis. J. Proteome Res. 6, 654–61. doi:10.1021/pr0604054

Tschaplinski, T.J., Standaert, R.F., Engle, N.L., Martin, M.Z., Sangha, A.K., Parks, J.M., Smith, J.C., Samuel, R., Jiang, N., Pu, Y., Ragauskas, A.J., Hamilton, C.Y., Fu, C., Wang, Z.-Y., Davison, B.H., Dixon, R.A., Mielenz, J.R., 2012. Down-regulation of the caffeic acid O-methyltransferase gene in switchgrass reveals a novel monolignol analog. Biotechnol. Biofuels. 5, 71. doi:10.1186/1754-6834-5-71

Wang, W., Papov, V. V., Minakawa, N., Matsuda, A., Biemann, K., Hedstrom, L., 1996. Inactivation of Inosine 5’-Monophosphate Dehydrogenase by the Antiviral Agent 5-Ethynyl-1-β-d-Ribofuranosylimidazole-4-Carboxamide 5’-Monophosphate^†^. Biochemistry 35, 95–101. doi:10.1021/bi951499q

Yi, X., Gu, H., Gao, Q., Liu, Z.L., Bao, J., 2015. Transcriptome analysis of Zymomonas mobilis ZM4 reveals mechanisms of tolerance and detoxification of phenolic aldehyde inhibitors from lignocellulose pretreatment. Biotechnol. Biofuels. 8, 153. doi:10.1186/s13068-015-0333-9

Zimmermann, L., Stephens, A., Nam, S.-Z., Rau, D., Kübler, J., Lozajic, M., Gabler, F., Söding, J., Lupas, A.N., Alva, V., 2017. A Completely Reimplemented MPI Bioinformatics Toolkit with a New HHpred Server at its Core. J. Mol. Biol. doi:10.1016/J.JMB.2017.12.007

